# Genetics, and not diet, drives DNA methylation in the water strider *Microvelia longipes*

**DOI:** 10.64898/2026.08.25.746969

**Authors:** Mirjam Urb, Séverine Viala, Juliette Mendes, Abderrahman Khila

## Abstract

Phenotypic plasticity, the ability of a single genotype to produce alternative phenotypes in response to environmental cues, is a key driver of evolutionary change. In the water strider *Microvelia longipes*, males display remarkable continuous variation in hindleg length, a sexually selected trait used as a weapon in male–male contests for access to females. To determine whether DNA methylation mediates this environmentally induced phenotypic variation, we used three inbred lines of *M. longipes* showing differences in mean hindleg length, body size, and allometric coefficients. Whole-genome bisulfite sequencing (WGBS) on adult males and females from all lines determined that about 12% of the 12,684,876 CpG sites found in the genome were methylated. This global level of DNA methylation is among the highest reported in insects. DNA methylation was predominantly concentrated within or near gene bodies (77% of methylated CpGs), consistent with patterns observed in other insects. Unsupervised clustering and principal component analyses revealed that methylation patterns differed significantly between genetic lines but showed minimal differences between sexes, indicating a strong genetic influence. Most surprisingly, nutritional treatment followed by leg-specific WGBS failed to identify any change in DNA methylation despite nutrition having a pronounced effect on leg length. These results show that in *M. longipes*, DNA methylation patterns are largely stable across nutritional treatments and primarily determined by genetic background. This challenges the common assumption that DNA methylation universally mediates environmentally induced phenotypic plasticity and suggests that other epigenetic mechanisms, such as histone modifications or non-coding RNAs, may play a more direct role in regulating continuous plastic traits. Our study underscores the complexity of epigenetic regulation and highlights the need for broader investigation of molecular pathways to fully understand the molecular basis of phenotypic variation in natural populations.

## Introduction

Phenotypic plasticity, defined as the ability of a single genotype to produce alternative phenotypes in response to environmental variation, is widely recognized as a key driver of evolutionary change across the tree of life (West-Eberhard 2003). Environmentally induced phenotypic changes can be discrete or continuous. In the case of discrete change, populations exhibit alternative morphs that are recognizable through clear quantitative (e.g., size) or qualitative (e.g., shape, developmental time, color) differences. Examples of plastically induced alternative morphs are common in frogs, insects, worms, crustaceans, vertebrates, plants and many other lineages (West-Eberhard 2003, Pfennig, Wund et al. 2010). In natural populations of the tadpoles of the spade-foot toad *Scaphiopus multiplicatus* for example, slow developing omnivorous and fast developing carnivorous tadpole morphs are induced environmentally, and the proportion of the carnivorous morph is determined by the abundance of shrimps, which represent their main prey (Pfennig 1990). In many insects, winged or wingless morphs develop in response to various environmental cues, such as nutrition (Huang, Zhang et al. 2023), photoperiod (Gudmunds, Narayanan et al. 2022) and habitat quality or population density (Fairbairn and King 2008). In the nematode *Pristionchus pacificus*, a predatory morph, which is an alternative to bacteria feeding morph, develops teeth-like structures that allow it to prey on other worm species (Bento, Ogawa et al. 2010). Other examples of discrete environmentally induced phenotypes include alternative color morphs in insects and fishes (Endler 1980, Tanaka 2004), defensive armors (Nagano and Doi 2020), or sexually-selected ornaments and weapons, such as plumage type in ruffs (Lank, Smith et al. 1995), horns in beetles (Emlen 1994), or side blotches in some lizards (Micheletti, Parra et al. 2012).

The second effect of phenotypic plasticity manifests in the environmental induction of continuous phenotypic variation. This is a far more common phenomenon as most organisms exhibit a certain degree of responsiveness to environmental change. Continuous plastic traits are typically considered complex, and the extent of their variation along a spectrum can differ dramatically among traits and lineages (Nijhout 2003). Genetically, these traits are quantitative in that they are shaped during development through the additive effect of a large number of loci and complex genetic and gene-by-environment interactions (Lynch and Walsh 1998, Nijhout 2003). Exaggerated sexually selected traits represent some of the most variable types of continuous plastic traits and are often found in males (Emlen 2008, Casasa, Schwab et al. 2017, Toubiana and Khila 2019, Miller, Cram et al. 2026).

Recent advances in our understanding of the mechanistic underpinnings of the effect of the environment on phenotypic variation have primarily focused on the study of discrete alternative phenotypes (West-Eberhard 2003, Sommer 2020). This bias is possibly due to the unambiguous nature of the alternative morphs, making it more practical in studies seeking to determine readout upon environmental perturbations (Sommer 2020). In contrast, continuous variation in plastic traits comes with the difficulty of disentangling the contribution of environmental variation from that of genetic variation (Sommer 2020). For example, studies in insects implicated the role of conserved components of the growth pathways in mediating the switch between alternative wing morphs. Interestingly, different environmental cues seem to induce alternative morphs through different growth pathways (Xu, Xue et al. 2015, Gudmunds, Narayanan et al. 2022). In plant hoppers, the switch between long and short wing morphs is induced by nutrition and mediated through the insulin pathway (Xu, Xue et al. 2015). In some water striders, where wing polymorphism is determined by photoperiod, the insulin pathway is not involved (Gudmunds, Narayanan et al. 2022) and the switch between long and short wing morphs is rather mediated through the Hipo pathway (Gudmunds, Palahí et al. 2024). In the worm *P. pacificus*, a gene named *eud-1* has been implicated in mediating the environmentally-induced switch between predatory and non-predatory morphs (Ragsdale, Müller et al. 2013). These studies in discrete plastic phenotypes have been important to better understand the mechanistic underpinnings of plastic responses, but the extent to which the gleaned conclusions apply to continuous traits is limited. The recent technological advances combined with experimental tractability of newly developed study models call for a more significant inclusion of continuously plastic traits in studies of the molecular mechanisms underlying environmentally induced variation.

Here, we investigate a continuously variable sexually selected trait in the water strider *Microvelia longipes* (Andersen 1982, Toubiana and Khila 2019). Males of this species exhibit extreme variation in hindleg length, a trait used as a weapon during male–male contests for access to females (Toubiana and Khila 2019, Toubiana, Armisen et al. 2021). In contrast, females display little variation in hindleg length and possess significantly shorter legs than males (Toubiana and Khila 2019). In males, hindleg size exhibits pronounced hyperallometry, with an allometric coefficient exceeding 3, among the highest reported in arthropods (Toubiana and Khila 2019).

*Microvelia longipes* is widely distributed across tropical regions where it inhabits small ephemeral water bodies such as rain puddles, which are prone to rapid fluctuations in water level. Females preferentially oviposit on small floating debris (e.g., fragments of twigs or dead leaves). These sites are guarded by dominant males, which produce signals by rhythmically striking the water surface with their genitalia, generating ripple-based calls to attract females (Toubiana and Khila 2019). Dominant males are frequently challenged by rivals in aggressive contests that involve striking with their elongated hindlegs in attempts to displace the resident male from the egg-laying site. These contests are typically won by larger males, underscoring the role of sexual selection in driving the evolution of exaggerated hindleg length in *M. longipes* (Toubiana and Khila 2019). Furthermore, reaction norm experiments revealed that growth of the hind legs of the males in particular is sensitive to nutritional treatment (Toubiana and Khila 2019), suggesting that nutritional treatment could induce growth differences through changes in DNA methylation and gene expression.

It has long been hypothesized that phenotypic variation is induced through variation in gene expression during development (Waterland and Jirtle 2003, Gilbert 2005). In the case of environmentally-induced phenotypic variation, changes in gene expression are thought to be induced through epigenetic modifications. These are chemical modifications of DNA bases that don’t alter the genomic sequence (Jablonka and Raz 2009, Glastad, Hunt et al. 2019). A primary mechanism that chemically alters the genome without changing DNA sequence is DNA methylation (Glastad, Hunt et al. 2019). Studies of the link between DNA methylation and phenotypic variation in non-conventional models often face difficulties of noise due to DNA polymorphism found in natural populations. Here, we used inbred lines of *M. longipes* to control for the effect of DNA polymorphism on phenotypic variation. First, we performed whole-genome bisulfite sequencing on whole adult males and females of three inbred lines which vary in their mean hindleg length and body size. This allowed us to determine the general rate and genomic pattern of DNA methylation for the first time in this species and the differences in DNA methylation sites between sexes and between lines. We then tested more specifically the effect of DNA methylation patterns on the growth of hindlegs of one of these inbred lines treated with poor or rich diet. Our data reveal the extent of differential methylation between lines with large and small legs, as well as between the sexes, and indicate that these patterns depend more on genetic background than nutritional status or sex.

## Results and Discussion

### Inbred lines of *M. longipes* show significant differences in leg length and body size

In this study, we used three inbred lines of *M. longipes* that exhibit differences in mean body size and hind leg length (**Fig. 1**). These lines are referred to as LH (long legs, high slope), SH (short legs, high slope), and LL (long legs, low slope) (**Fig. 1a-b**), although the slopes are not statistically significant (Males: LH vs LL *p*=0.1793, LH vs SH *p*=0.8248, LL vs SH *p*=0.7764; Females: LH vs LL *p*=0.2788, LH vs SH *p*=0.2963. LL vs SH *p*=0.0154. The most recent phenotypic data for these lines is presented in **Fig. 1**. The LH line consistently displays longer legs and large body size; the SH line exhibits smaller body; and the LL line is intermediate in both traits for males and females (Male hind tibia: LH vs LL *p*=5.2e-15, LH vs SH *p*<2.22e-16, LL vs SH *p*=0.034; Female hind tibia: LH vs LL *p*=0.14, LH vs SH *p*=2.3e-11, LL vs SH *p*=1.8e-09; Male body: LH vs LL *p*=2.9e-09, LH vs SH *p*<2.22e-16, LL vs SH *p*=5.7e-11; Female body: LH vs LL *p*=0.022, LH vs SH *p*<2.22e-16, LL vs SH *p*=1.2e-15; **Fig. 1c-f**). We, therefore, used these lines to determine the genomic patterns of DNA methylation and the differences in these patterns across lines with different phenotypes.

**Figure 1:**
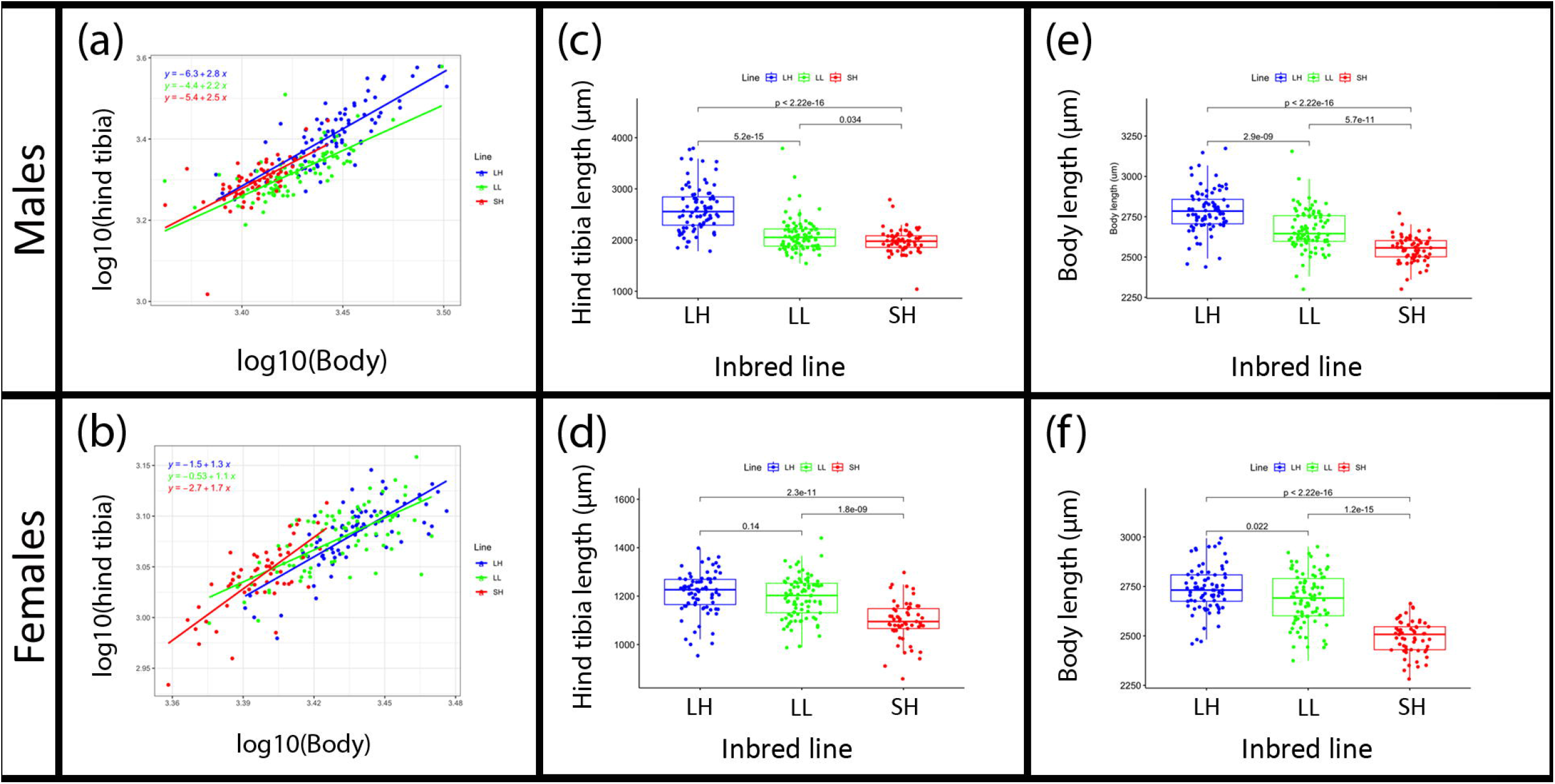
Phenotyping of the three inbred lines used in this study, at the adult stage. (**a-b**) Scaling relationships between hind tibia and body in the males (**a**) and females (**b**) of the three lines. (**c-f**) Boxplots showing differences in leg length (**c-d**) and body length (**e-f**) in males and females.

### High levels of CpG DNA methylation in *M. longipes*

DNA methylation has been proposed as one of the key epigenetic mechanisms underlying phenotypic plasticity in a wide range of organisms (Ibañez, Masuelli et al. 2021, Duncan, Cunningham et al. 2022, Venney, Anastasiadi et al. 2023, Bogan and Yi 2024). To begin to investigate whether the observed extreme growth variation in *M. longipes* is mediated by changes in DNA methylation, we first analyzed methylation patterns across the genome in the adults of all three lines. The latest *M. longipes* genome assembly spans approximately 670 megabases, with 90% of the sequence contained within the 13 largest scaffolds (Toubiana, Armisen et al. 2021). Our analysis identified 12,684,876 CpG sites, 14,365,104 CHG sites, and 55,049,107 CHH sites per strand (where H represents A, C, or T). Automatic genome annotation (Toubiana, Armisen et al. 2021), confirmed by manual annotation revealed the presence of DNMT1 as a single representative of DNA methyl transferases, but failed to detect DNMT2 and DNMT3 in *M. longipes* (Toubiana, Armisen et al. 2021). Although DNMT1 is primarily known for its role in maintaining CpG DNA methylation across cell divisions (Hermann, Goyal et al. 2004), it can also play a role in de novo methylation in some biological contexts such as transposable elements (Haggerty, Kretzmer et al. 2021). Additionally, DNMT1 is known to influence phenotypic responses to environmental changes in some insects (Tang, Zhang et al. 2024), further reinforcing the hypothesis that DNA methylation could contribute to male phenotypic variation in *M. longipes*.

Whole-genome bisulfite sequencing (WGBS) of adult individuals from the three inbred lines revealed that about 12% of CpG sites were methylated, with a highly consistent pattern across all lines (**Fig. 2a**). This level of CpG methylation ranks among the highest reported in insects (Bewick, Vogel et al. 2017). In contrast, CHG and CHH sites showed methylation levels below 1% (0.3%) across the genome (**Fig. 2a**). These findings indicate that the rate of DNA methylation in *M. longipes* is quite high among insects, with methylation largely confined to CpG sites.

**Figure 2:**
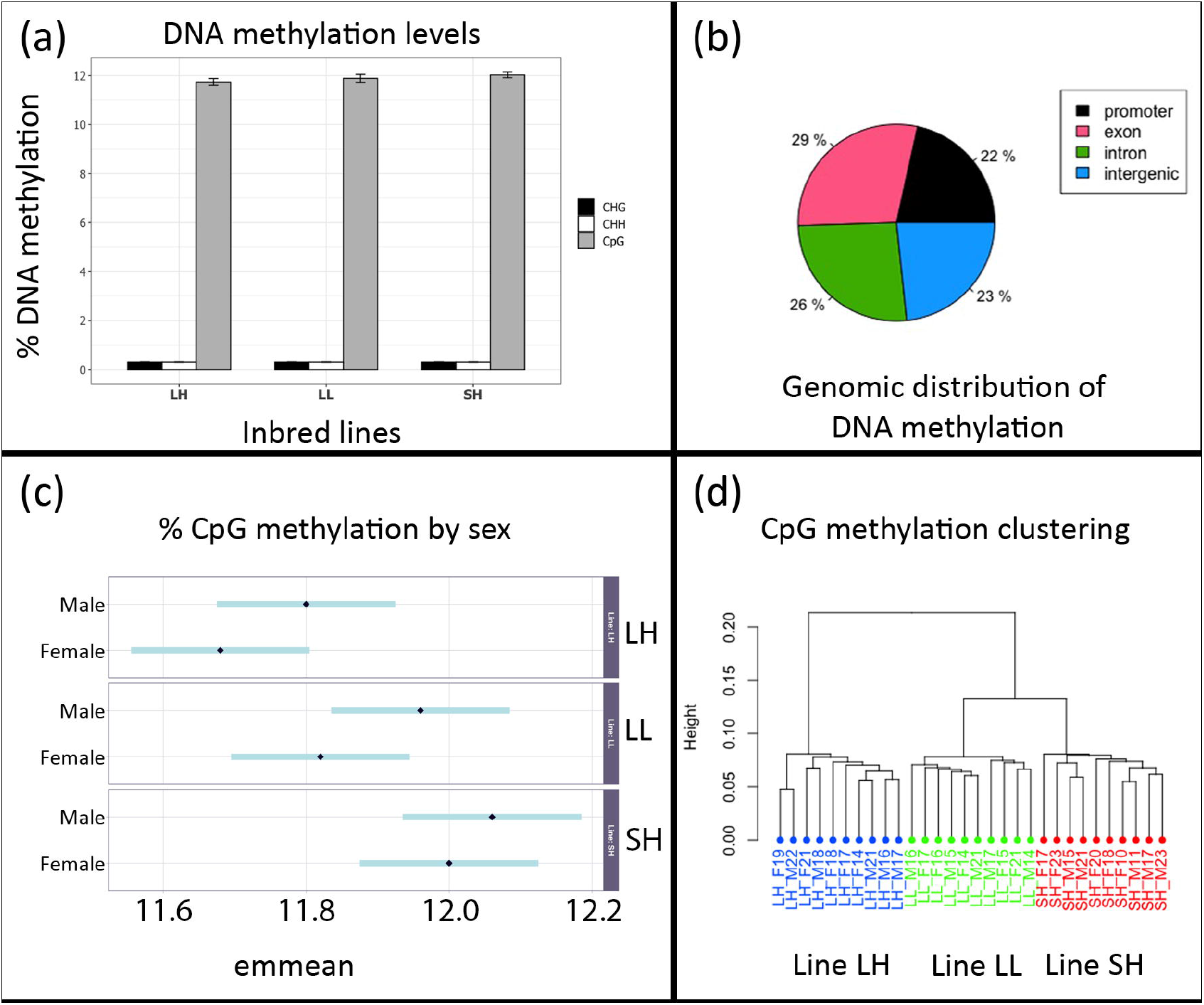
Genome-wide DNA methylation analyses in *M. longipes*. (**a**) CHG, CHH and CpG DNA methylation levels across the three lines. (**b**) Genomic distribution of CpG DNA methylation patterns in *M. longipes*. (**c**) Sex differences in global levels of CpG DNA methylation in *M. longipes* by line. (**d**) Unsupervised clustering of DNA methylation comparing lines and sexes.

A detailed genome-wide analysis revealed that DNA methylation in *M. longipes* is primarily concentrated within or near gene bodies (**Fig. 2b**). Only 23% of methylated CpG sites were located in intergenic regions (**Fig. 2b**), while the remaining 77% of the sites were distributed across exons (29%), introns (26%), and promoter regions (22%) (**Fig. 2b**). This means that more than three-quarters of methylated CpGs are concentrated within or near protein coding sequences, which represent a small fraction of the genome. These data are consistent with previous findings in insects, and suggest that methylation may play a role in the regulation of gene expression either through transcriptional or translational control (Neri, Rapelli et al. 2017, Shahib and Rastegar 2026).

A comparative analysis between males and females showed a slight, but statistically nonsignificant (LH: *p*=0.1739, LL: *p*=0.1151, SH: *p*=0.4903), difference in overall CpG methylation levels, with a trend toward higher methylation in males (**Fig. 2c**). Overall, these results indicate that global DNA methylation levels and genomic distribution patterns in *M. longipes* are largely stable across inbred lines and between the sexes, with a pronounced enrichment in genic regions.

### Methylation patterns vary more between lines than between sexes

Unsupervised clustering analysis of DNA methylation levels grouped the samples according to their genetic line (**Fig. 2d**). The LH line, characterized by males with larger leg and body size, was distinctly separated from the other two lines, which clustered closely together (**Fig. 2d**). This separation is reflected in the number of differentially methylated CpG sites between lines: 2,485 between LH and SH, 1,520 between LH and LL, and 802 between LL and SH **(Table 1)**. These numbers increased significantly when the cutoff was relaxed to 10% (**Fig. 3**; **Table 1**), indicating that more phenotypically distinct lines exhibit greater differences in methylation patterns. The CpG methylation differences between lines are equally represented by hyper- and hypomethylation (**Fig. 3; Table 1**).

**Table 1:**
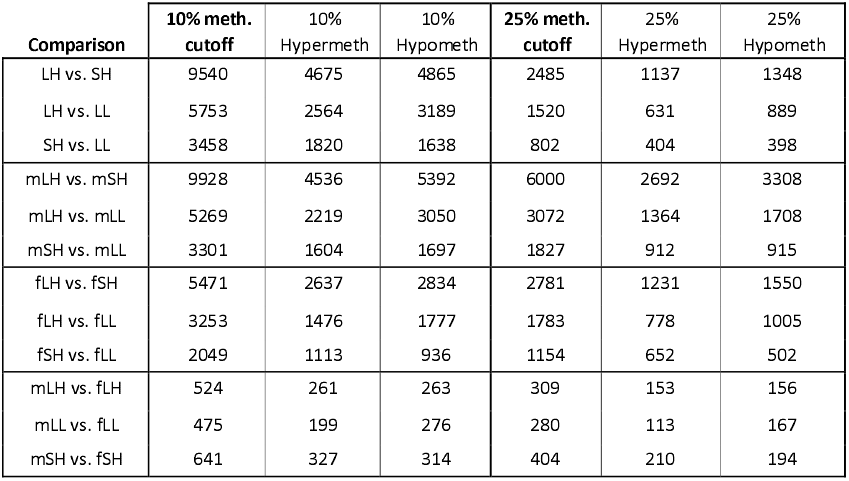
Number of differentially methylated sites between lines and sexes. m: Males; f: Females. LH: Long legs/high slope line; LL: Long legs/low slope line; SH: Sort legs/high slope line.

**Figure 3:**
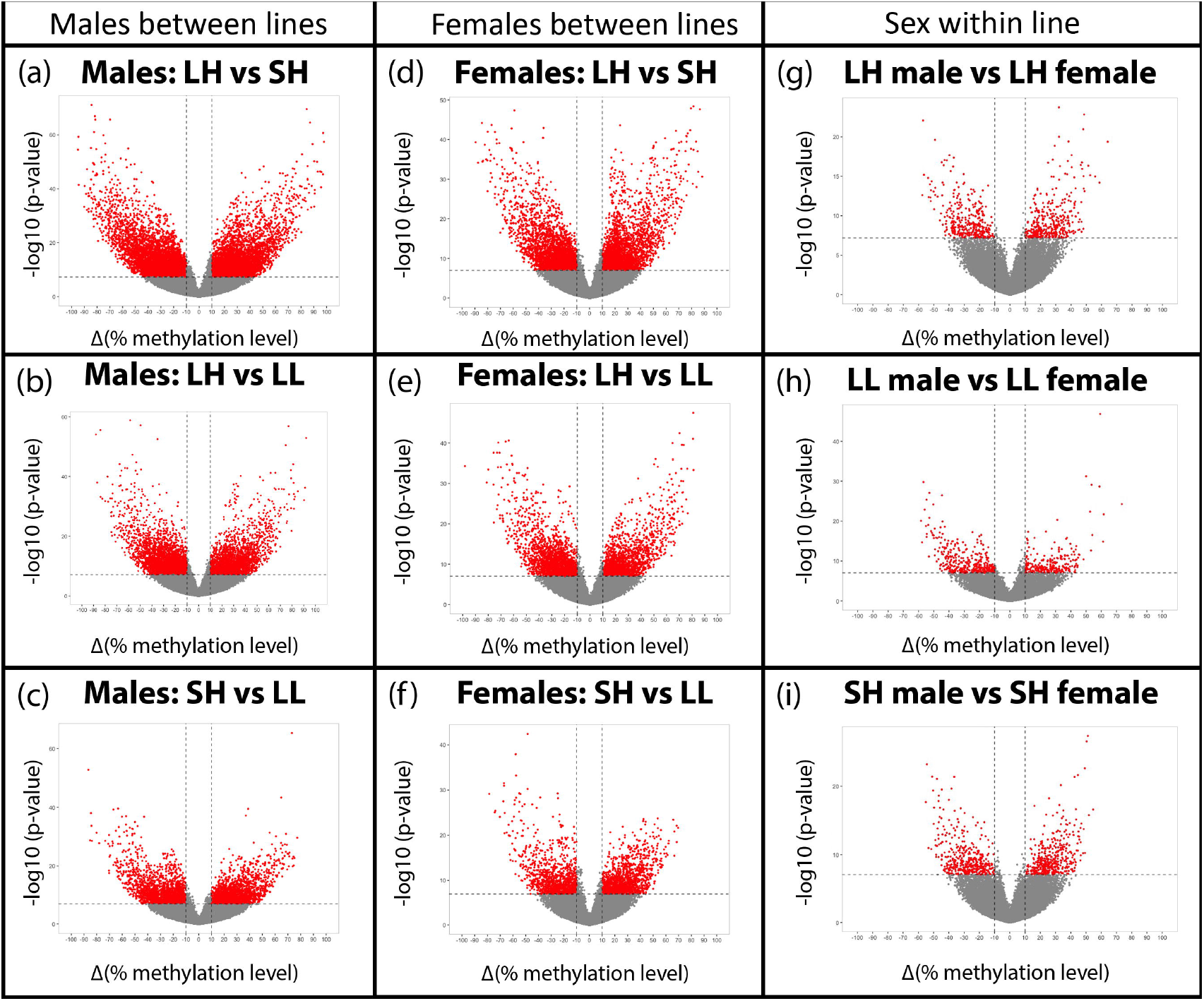
Differential CpG methylation across lines and between the sexes. (**a-c**) Differential methylation between males of the different lines. (**d-f**) Differential methylation between females of the different lines. (**g-h**) Differential methylation between males and females of the same line. The points in red represent the significant sites with >=10% differential methylation.

When comparing males (**Fig. 3a-c**) and females (**Fig. 3d-f**) separately across the three lines, the extent of DNA methylation mirrored the phenotypic differences observed between lines (**Fig. 1**). For example, the LH line, where males are the largest on average, differed from the SH line, where males are the smallest, by some 6,000 methylated sites (25% cutoff), with about 45% being hyper- and the other 55% hypomethylated (**Fig. 3; Table 1**). In other comparisons, methylation differences were intermediate between males of LH and LL lines and lower between males of LL and SH lines (**Table 1**). In females, a similar trend in methylation differences was observed, but with about half the numbers of differentially methylated sites than in males (**Table 1**). Thus, DNA methylation differences align with phenotypic variation, particularly in males (**Fig. 1**).

Notably, this analysis failed to cluster males and females separately within lines based on DNA methylation differences (**Fig. 2d**). The number of differentially methylated sites between lines is about an order of magnitude higher than the differences between sexes within a given line (**Fig. 3**; **Table 1**). Principal component and unsupervised clustering analyses performed on each line separately also failed to group samples by sex (**Supplementary Fig. S1**). Therefore, global DNA methylation patterns are consistent among individuals of the same line, regardless of sex, but differ significantly between lines. This indicates a stronger influence of genetic background than sex on DNA methylation patterns in this species.

Given the stability of global DNA methylation patterns within lines, we investigated whether methylation levels differ between males and females at a finer, site-specific scale. Scatterplot analysis identified specific sites that were differentially methylated between sexes within each line (**Table 1**; **Fig. 3g–i**). All lines showed consistent numbers of DNA methylation differences between sexes, ranging from 280 to 400 (cut off 25%) sites, with roughly half being hypo- and the other half hyper-methylated (**Table 1**). This number increased when applying the more relaxed 10% cutoff (**Table 1**). Surprisingly, very few differentially methylated sites were common to all lines (**Fig. 4a**), and no sex-biased sites were shared among all three lines (**Fig. 4b**), further emphasizing the strong effect of genetic background on DNA methylation profiles. A genome-wide analysis revealed that methylation differences were spread across the 13 largest chromosomes, which represent 90% of the genome (Toubiana, Armisen et al. 2021) (**Supplementary Fig. S2; Table 1**). Peaks of methylation patterns often comprised both hypo- and hypermethylated sites, both passing statistical significance (**Supplementary Fig. S2**), suggesting that these genomic regions may be subject to variation in DNA methylation states.

**Figure 4:**
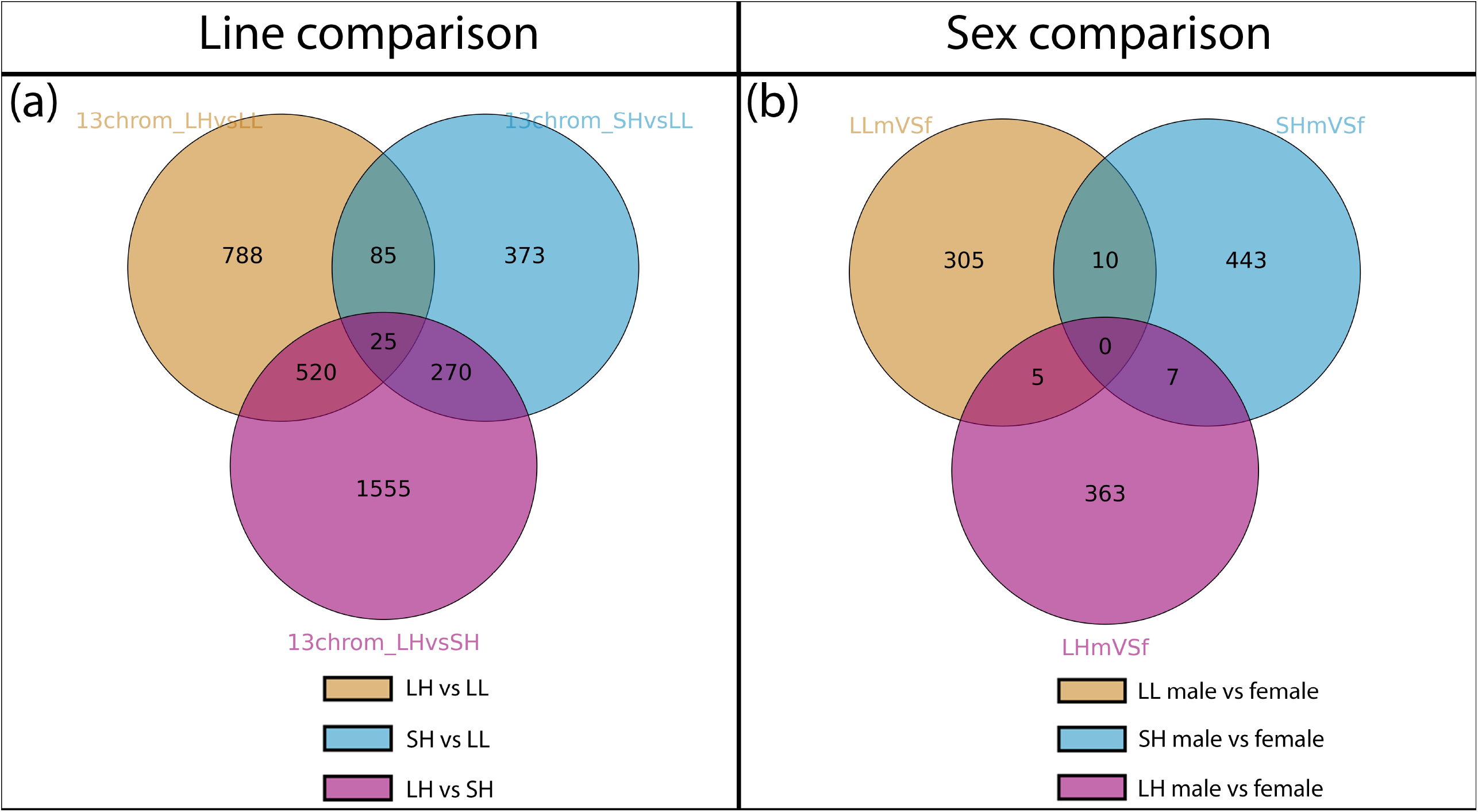
Common DNA methylation patterns between lines and sexes. **(a)** Only 25 differentially methylated sites are shared between all three lines **(b)** No differentially methylated sites between the sexes are common to all three lines. Analysis was performed based on the 13 largest scaffolds containing 90% of *M. longipes* genome.

### DNA methylation in *M. longipes* does not respond to nutritional condition

Next, we wanted to test whether the growth variation of *M. longipes* males we observe following nutritional treatment is mediated through changes in DNA methylation states (Toubiana and Khila 2019). We performed this test specifically on the hindlegs using the LH inbred line where males have the longest hindlegs among the lines (**Fig. 1**). Second instar nymphs were raised on either poor or rich diet till they reached the fifth instar (**Supplementary Fig. S3**). The effect of diet on leg length was significantly pronounced on these samples (p<2.22e-16; **Supplementary Fig. S3**), confirming previous findings that leg length is heavily influenced by nutritional intake (Toubiana and Khila 2019). Second day 5^th^ instar nymphs were dissected and DNA of pools of 40 legs per replicate was extracted to conduct whole-genome bisulfite sequencing. General hierarchical clustering, including all leg samples and nutritional conditions, looking for dissimilarities in global methylation per base across samples failed to detect any clear pattern based on nutritional treatment or sex (**Supplementary Fig. 4a**). This observation was confirmed in PCA analysis where all samples clustered together, except of two replicates of forelegs from animals raised on poor diet (**Supplementary Fig. 4b**). Binary comparisons between forelegs and hind legs in males and females of individuals raised on rich or poor diet also failed to cluster samples by diet (**Fig. 5**). The lack of genes showing differences in DNA methylation status in response to nutritional treatment is surprising, particularly given that the experiment is well controlled, replicated and the depth of sequencing is above average. We conclude that DNA methylation in *M. longipes*, despite its high levels, is robust to nutritional changes.

**Figure 5:**
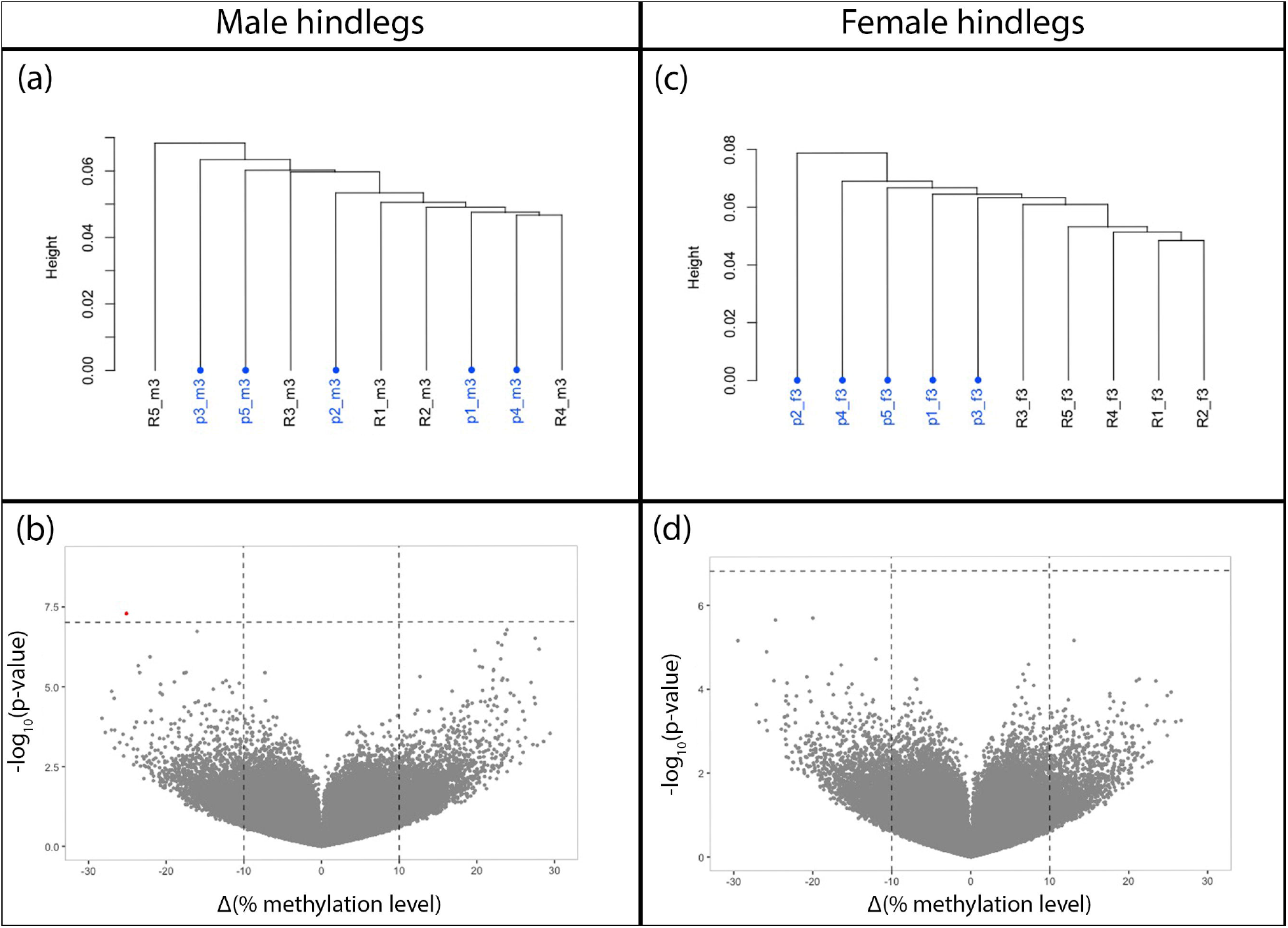
Hierarchical clustering (**a** and **c**) and 10% differential CpG methylation (**b** and **d**) between the hindlegs of males and females raised on poor or rich diet. R: rich diet; P: poor diet; numbers indicate replicates.

Our findings reveal that *M. longipes* exhibits high levels of CpG methylation, primarily concentrated within or near gene bodies. The methylation landscape in *M. longipes* is remarkably consistent across inbred lines and sexes, with only subtle differences observed between males and females. Surprisingly, CpG methylation patterns seem to be highly robust to nutritional variation, even though this environmental cue has a profound effect on the final phenotypes in the males of *M. longipes*. Another striking result is that methylation patterns differ more between lines than between sexes, indicating that genetic background has a stronger influence on methylation than sex-specific or environmental factors. These findings suggest that DNA methylation in *M. longipes* is primarily influenced by genetics, and that methylation patterns may potentially be passed across generations. However, the stability and functional significance of these inherited marks on the phenotype of each line remains to be tested. Further research could explore whether these methylation patterns are adaptive or constitute simple passive modifications with little effect on the final phenotype.

One of the most unexpected findings of this study is that DNA methylation in *M. longipes* does not respond to nutritional treatment. This result is particularly surprising given the well documented role of nutrition in inducing phenotypic plasticity in other species, such as the insulin-mediated wing polymorphism in plant hoppers or the photoperiod-induced wing dimorphism in water striders (Xu, Xue et al. 2015, Gudmunds, Narayanan et al. 2022). In *M. longipes*, where hindleg length is a continuously variable trait subject to sexual selection (Toubiana and Khila 2019), one might expect nutritional conditions to influence methylation patterns, thereby modulating leg growth. However, our data suggest that DNA methylation in this species is robust to nutritional changes, at least under the conditions tested. This lack of responsiveness could imply that the extreme variation in hindleg length is primarily driven by other epigenetic mechanisms, such as histone modifications (Glastad, Hunt et al. 2019) or non-coding RNAs (Lee, Feinbaum et al. 1993). Future studies could explore these alternative pathways to better understand the molecular basis of phenotypic plasticity in *M. longipes*.

In summary, this study provides a comprehensive analysis of DNA methylation in *M. longipes*, revealing that methylation patterns are strongly influenced by genetic background but surprisingly unresponsive to nutritional conditions. These findings underscore the complexity of phenotypic plasticity and the need for further research to disentangle the relative contributions of genetic, epigenetic, and environmental factors in shaping continuous plastic traits. By expanding our understanding of these mechanisms, we can gain deeper insights into the evolutionary processes that generate and maintain phenotypic diversity in natural populations.

## Material and Methods

### Animals

The original *Microvelia longipes* population was collected from French Guyana in 2013. From this population three isogenic lines were established from 20 generations of sibling-sibling inbreeding (Toubiana and Khila 2019), and they have been maintained ever since. These lines are referred to as LH (long legs, high slope), SH (short legs, high slope) and LL (long legs, low slope). The bugs were housed in the laboratory at 28-29°C with 50-60% humidity and 14 hours of daylight in water buckets, and were fed crickets daily.

### Leg and body measurements

To assess the leg and body length of the current isogenic lines, images were taken with Nikon D7200 camera with AF-S Micro-NIKKOR 105 mm lens, and measured using ImageJ v. 2.14.0/1.54f. Tibia of the hindleg was used as a proxy for the total leg length (Toubiana and Khila 2019). Allometry and boxplots were generated with ggplot2 v3.5.1 (Wickham 2009). The statistical analyses were performed with R packages emmeans v2.0.4 (Lenth and Piaskowski 2026) and ggpubr v0.60 (Kassambara 2026). P-values were calculated using pairwise Tukey test and Kruskal-Wallis test, respectively.

### Sample collection, genomic DNA extraction and whole-genome bisulfite sequencing

Adult males and females from the stock population of all three isogenic lines were isolated at 5^th^ instar to obtain virgin adults. The freshly emerged adults were starved for 48 h to prevent cricket contamination, and were stored at -80°C until genomic DNA extraction. Of note, growth in insects stops at the adult stage and no change in size occurs after.

Genomic DNA from five replicates of each sex of the three isogenic lines was extracted from whole individuals with NucleoSpin-Tissue kit (Macherey-Nagel) according to manufacturer instructions. The samples were sent to SNP&SEQ Technology Platform in Uppsala for Whole Bisulfite Sequencing (150 cycles of paired-end sequencing in 1 lane of a S4 flowcell using NovaSeq 6000 system and v1.5 sequencing chemistry; Illumina Inc.). Sequencing libraries were prepared from 200 ng of DNA using the SPLAT method according to the in-house protocol (Raine, Manlig et al. 2017).

### DNA methylation analyses

Sequencing quality of both raw and trimmed reads was evaluated using FastQC v0.11.9. FastQ Screen v0.23.1 in bisulfite mode was used to detect possible contaminations against a set of sequence databases (Wingett and Andrews 2018). FastQC and FastQ Screen results were summarized with MultiQC v1.12 (Ewels, Magnusson et al. 2016). Trimming was conducted using TrimGalore v0.6.6 (Krueger 2015) that requires the software Cutadapt (Martin 2011). The trimmed sequences were aligned to bisulfite-converted and indexed *M. longipes* reference genome (Toubiana, Armisen et al. 2021) in paired-end and non-directional mode using Bismark v0.23.1 (Krueger and Andrews 2011) with Bowtie2 v2.4.4 (Langmead and Salzberg 2012). Before methylation calling with Bismark, the sequences were PCR deduplicated, name sorted, mate fixed, position sorted, and duplication marked and removed with samtools v1.17 (Li, Handsaker et al. 2009). DNA methylation percentage was calculated by dividing methylated C’s with the sum of methylated and unmethylated C’s times 100.

All subsequent DNA methylation analyses and filtering were performed in RStudio v. 4.2.2 (Team 2020) using methylKit v1.24.0 (Akalin, Kormaksson et al. 2012). Raw data was used to investigate the clustering between samples in CpG context with both unhierarchical clustering and principal component analysis. From here on, pairwise comparisons between or within isogenic lines were only performed. Samples were filtered by discarding bases with a coverage below 10x and over the 99.9th percentile to avoid PCR biases. Next, the filtered data was destranded to merge CpG sites from both strands, and only CpGs that were shared between all five replicates were included. Differentially methylated CpG sites were calculated using Bonferroni adjustment, and 10% and 25% percent methylation differences between two conditions were used in the graphs and tables.

Volcano plots were generated with ggplot2 v3.5.1 (Wickham 2009) and Manhattan plots with ggman v0.99.0 (Turner 2018). To look for overlaps between differentially methylated sites between different comparisons, bedtools v2.30.0 (Quinlan and Hall 2010) was used, and Venn diagrams created with Intervene v0.6.4 (Khan and Mathelier 2017).

### Nutritional experiment

This experiment focuses on the hindlegs of males, since this is the tissue that responds the most to changes in diet (Toubiana and Khila 2019). To investigate the potential nutritional effect on DNA methylation in the hind leg length, we exposed *M. longipes* LH line nymphs to either rich or poor diet. To standardize the age, first instar nymphs hatched within a 6-hour period from the stock population were used. The 1^st^ nymphal instars were fed the same rich diet consisting of fresh crickets (1 fresh cricket per 50 individuals) twice a day. When most nymphs reached 2nd instar, the nymphs were randomly divided into rich and poor conditions. The rich group received excess of fresh crickets twice a day with 1 fresh cricket per 20 individuals during 2^nd^ instar, and 1 fresh cricket per 10 individuals for the remaining of the experiment. The poor group received frozen cricket legs once a day. Second instar nymphs were fed 1 frozen leg per 20 individuals, and the remaining of the nymphal instars 1 frozen leg per 10 individuals.

To obtain leg-specific DNA from individuals treated with poor or rich diet, 5^th^ instar nymphs were stored at -80°C two days post-molting until all nymphs reached the 5^th^ instar. The individuals were photographed with VHX-7000 macroscope (Keyence), and tibia of hindlegs were measured with VHX-7000 software. Genomic DNA was extracted from the forelegs and hindlegs of both males and females for comparison. Five replicates of each condition (diet and sex) corresponding to a pool of either hindlegs or forelegs from 40 individuals with the longest or shortest legs corresponding to the most extreme effects of rich and poor diet, respectively (**Supplementary Fig. S3**). Sequencing was conducted at the SNP&SEQ Technology Platform in Uppsala. DNA methylation analyses were carried out as mentioned above with the exception of using CpG sites that were common between three or more replicates out of the five replicates (not all, as above) in the filtering step.

## Author contributions

M.U. and A.K. designed experiments; M.U. performed experiments; M.U., S.V. and J.M. did the measurements; M.U. analyzed data; A.K. and M.U. wrote the manuscript.

## Acknowledgements

The authors thank Rita Rebollo for discussion and advice on DNA methylation analyses and Pascale Roux for technical assistance. We thank the PSMN (Pôle Scientifique de Modélisation Numérique) of the ENS de Lyon and from GENCI/TGCC (grant A0110807662) for their support and computing resources. Sequencing was performed by the SNP&SEQ Technology Platform in Uppsala. The facility is part of the National Genomics Infrastructure (NGI) Sweden and Science for Life Laboratory. The SNP&SEQ Platform is also supported by the Swedish Research Council and the Knut and Alice Wallenberg Foundation. This work was supported by an FRM équipe grant EQU202103012573 and VR 146700220 grant from Swedish research council grant to AK.

