## Supplementary online information for "Genetics, and not diet, drives DNA methylation in the water strider *Microvelia longipes*"

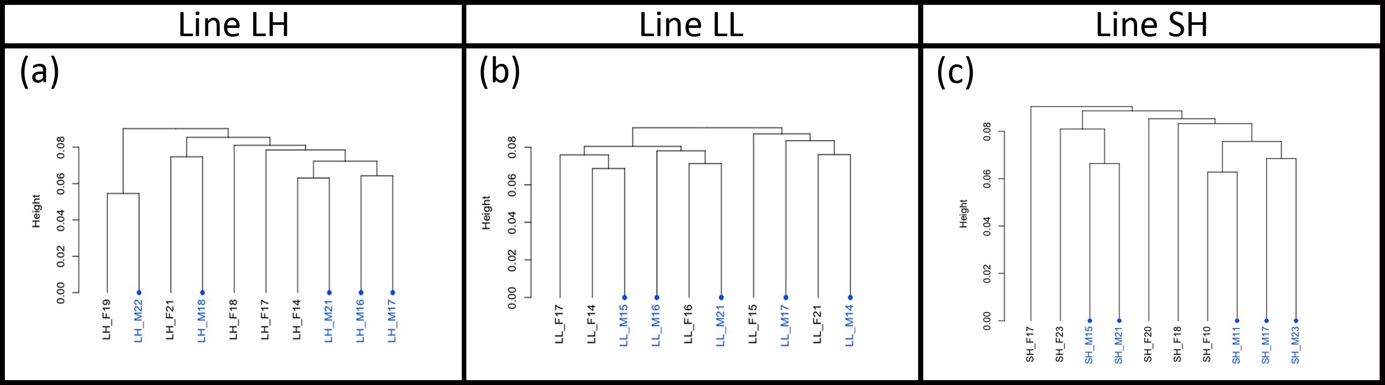

**Supplementary Figure S1:** Hierarchical clustering of CpG DNA methylation performed on five replicates from line LH (**a**), LL (**b**) or SH (**c**) fails to cluster samples by sex. M indicates males and F females, and the numbers represent different individuals.

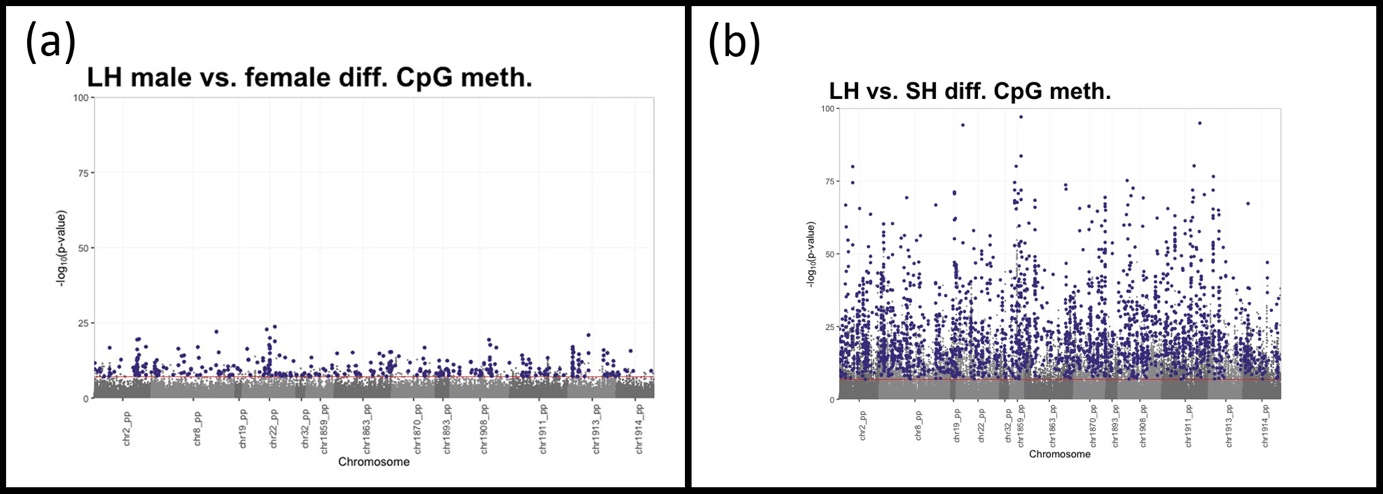

**Supplementary Figure S2**. Pattern of genomic distribution of significant differentially methylated CpG sites (25% cutoff) between males and females within the same line (**a**) and between two different lines (**b**). Bonferroni-corrected -log_10_(p) values are indicated by the red line, and significant CpG sites are represented by purple points.

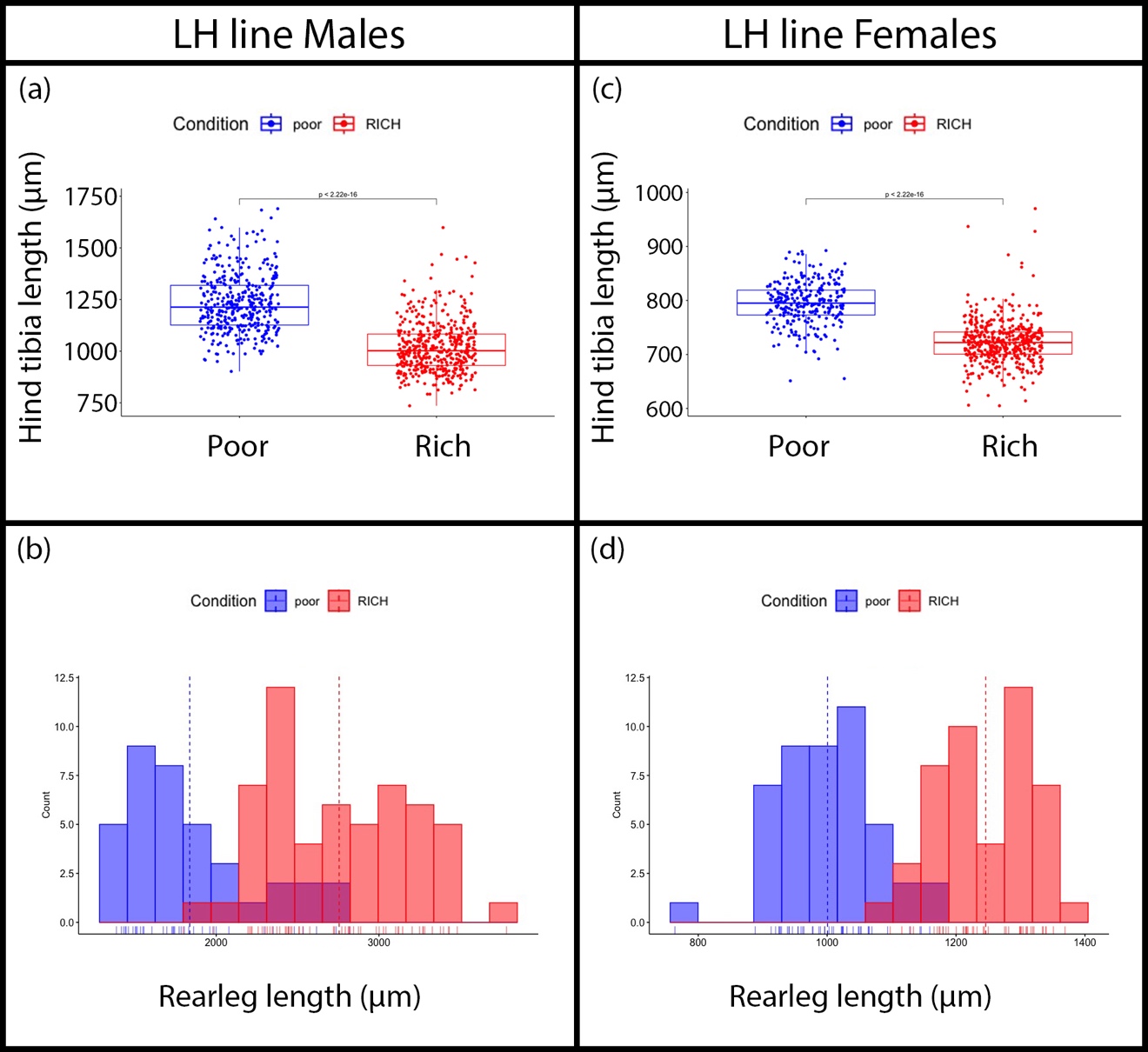

**Supplementary Figure 3:** Effect of poor (blue) and rich (red) diet on the growth of the hindlegs of *M. longipes* fifth instar nymphs. Note that this experiment was performed using the line LH. (**a-b**) Comparison of hindleg length in the males (**a**) and females (**b**) of fifth instar nymphs raised on rich or poor diet. (**c-d**) Hind leg length distribution of male (c) and female (d) fifth instar nymphs raised on poor (blue) or rich (red) diet.

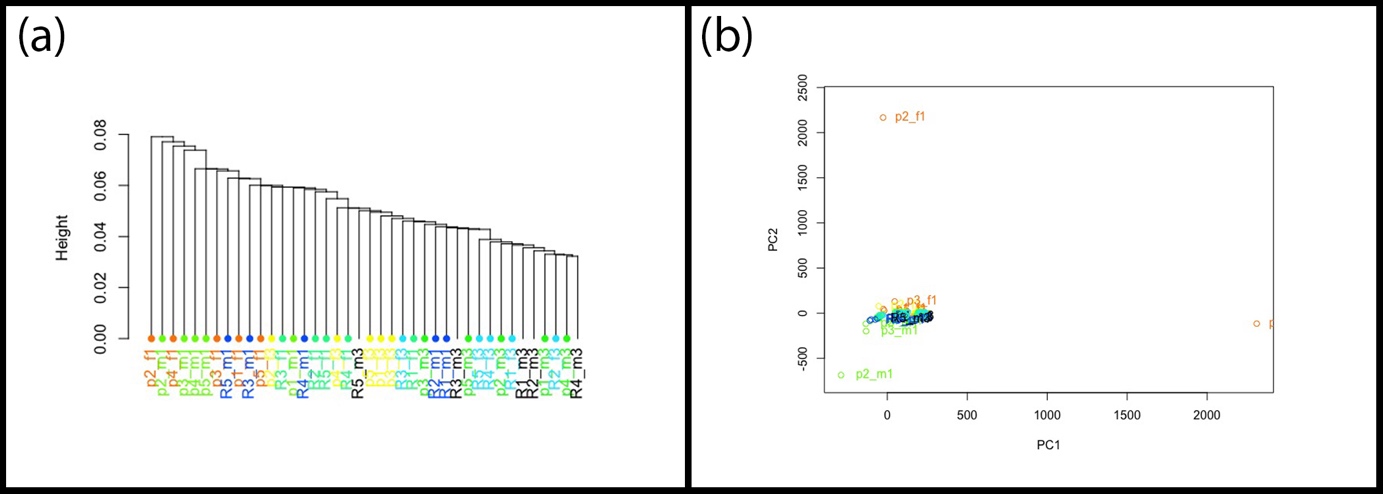

**Supplementary Figure 4:** Hierarchical clustering (**a**) and principal component analysis (**b**) fail to cluster samples by nutritional treatment. Brown: forelegs of females raised on poor diet; Turquoise: forelegs of females raised on rich diet; Light blue: hindlegs of females reared on rich diet; Yellow: hindlegs of females raised on poor diet; Dark blue: forelegs of males reared on rich diet; Black: hindlegs of males raised on rich diet; Light green: forelegs of males raised on poor diet; Dark green: hindlegs of males raised on poor diet. Numbers after rich (R) and poor (p) indicate replicates.
